# HOPS: Automated detection and authentication of pathogen DNA in archaeological remains

**DOI:** 10.1101/534198

**Authors:** Ron Hübler, Felix M. Key, Christina Warinner, Kirsten I. Bos, Johannes Krause, Alexander Herbig

## Abstract

High-throughput DNA sequencing enables large-scale metagenomic analyses of complex biological systems. Such analyses are not restricted to present day environmental or clinical samples, but can also be fruitfully applied to molecular data from archaeological remains (ancient DNA), and a focus on ancient bacteria can provide valuable information on the long-term evolutionary relationship between hosts and their pathogens. Here we present HOPS (**H**euristic **O**perations for **P**athogen **S**creening), an automated bacterial screening pipeline for ancient DNA sequence data that provides straightforward and reproducible information on species identification and authenticity. HOPS provides a versatile and fast pipeline for high-throughput screening of bacterial DNA from archaeological material to identify candidates for subsequent genomic-level analyses.

## Background

High-throughput DNA sequencing enables large-scale metagenomic analyses of environments and host tissues, providing an unprecedented understanding of life’s microbial diversity. Examples of coordinated efforts to quantify this diversity include the Human Microbiome Project (1), the Tara Ocean Project (2) and the Earth Microbiome Project (3). Metagenomic data from human archaeological remains (*e.g*. bones, teeth or dental calculus), which provide a window into the individuals’ metagenomic past, is a welcome addition to the wide landscape of microbial diversity now being revealed. While many ancient DNA (aDNA) studies focus on the analysis of human endogenous DNA isolated from ancient specimens (4–8), the co-recovered metagenomic aDNA can be queried to provide information related to endogenous microbial content at death, with applications ranging from characterizing the natural constituents of the microbiota to identifying systemic infectious diseases (9, 10).

Genomic-level investigations of ancient pathogens have provided valuable information about the evolution of *Yersinia pestis* (11–18), *Mycobacterium leprae* (19, 20), *Mycobacterium tuberculosis* (21, 22), pathogenic *Brucella* species (23, 24), *Salmonella enterica* (25, 26) and *Helicobacter pylori* (27), with others surely on the horizon. Notably, most studies to date have leveraged paleopathological evidence or historical context to pinpoint *a priori* involvement of a specific bacterial pathogen. However, the vast majority of infectious diseases do not lead to the formation of distinct and characteristic bone lesions, and most remains are found in contexts that lack clear associations with a particular disease. Consequently, studies of ancient pathogens must consider a long list of candidate microbes. Therefore, an automated computational screening tool that both detects and evaluates pathogen genetic signals in ancient metagenomic data is needed. Importantly, this tool should also be able to distinguish potential pathogens from the large and diverse microbial background typical of archaeological and other decomposed material, a consideration not typically required for tools developed for clinical applications.

To save computational time and effort, most available metagenomic profiling tools focus only on individual genes, such as the 16S rRNA gene used by QIIME (28), or panels of marker genes, such as those used by MetaPhlAn2 (29) and MIDAS (30), that are information-rich and highly species-specific. However, these genes make up only a small proportion of a bacterial genome (the 16S rRNA gene, for example, accounts for only ~0.2% of a bacterial genome), and if a pathogen is present at low abundance compared to host and environmental DNA, these genes are likely to be missed in routine metagenomic sequencing screens. As such, although these tools can be specific, they lack the sensitivity required for ancient pathogen screening from shallow metagenomic datasets. Screening techniques that accommodate queries of whole genomes are of clear benefit for archaeological studies (25). However, while some algorithms, such as Kraken (31), have been developed to query databases that contain thousands of complete reference genomes using k-mer matching, this approach does not produce the alignment information necessary to further evaluate species identification accuracy or authenticity.

In addition to taxonomic classification (32), it is also critical to distinguish ancient bacteria from modern contaminants (9, 10). Genuine aDNA, especially pathogen bacterial DNA, is usually only present in small amounts and can be distinguished from modern DNA contamination by applying an established set of authenticity criteria (9, 10), the most important of which is the assessment of DNA damage. In ancient DNA, cytosine deamination accumulates over time at DNA fragment termini (33, 34), thus leading to a specific pattern of nucleotide misincorporation. The evaluation of additional authenticity criteria such as edit distances and the distribution of mapped reads across the reference are also recommended to mitigate against database bias artifacts and to further validate taxonomic assignments (9, 10). While manual evaluation of species identification and aDNA authenticity using standalone tools might be feasible for a small sample set, it is impractical and too labour intensive to apply to the large sample sizes typical of recent ancient DNA investigations. The increased throughput of the ancient DNA field warrants an automated high-throughput solution for pathogen detection in metagenomic datasets. Successful ancient pathogen detection is reliant upon three criteria: (i) specificity of species-level detection against a diverse metagenomic background, (ii) high sensitivity that allows detection even with a weak signal when only trace amounts of species-specific DNA are present, and (iii) authentication of its ancient origin. However, no software currently exists that fulfills all requirements essential for reliable screening of metagenomic aDNA. Here we introduce HOPS (Heuristic Operations for Pathogen Screening), an automated computational pipeline that screens metagenomic aDNA data for the presence of bacterial pathogens and assesses their authenticity using established criteria. We test HOPS on experimental and simulated data and compare it to common metagenomic profiling tools designed for modern DNA analysis. We show that HOPS outperforms available tools, is highly specific and sensitive, and can perform reliable and reproducible taxonomic identification and authentication with as few as 50 reads.

## RESULTS

### HOPS Workflow

HOPS consists of three parts (Figure 1): i) a modified version of MALT (25, 35), which includes optional PCR duplicate removal and optional deamination pattern tolerance at the ends of reads; ii) The newly developed program MaltExtract, which provides statistics for the evaluation of species identification as well as aDNA authenticity criteria for a user-specified set of bacterial pathogens, with additional functionality to filter the aligned reads by various measures such as read length, sequence complexity or percent identity; and iii) a post-processing script that provides a summary overview for all samples and potential bacterial pathogens that have been identified.

**Figure 1.**
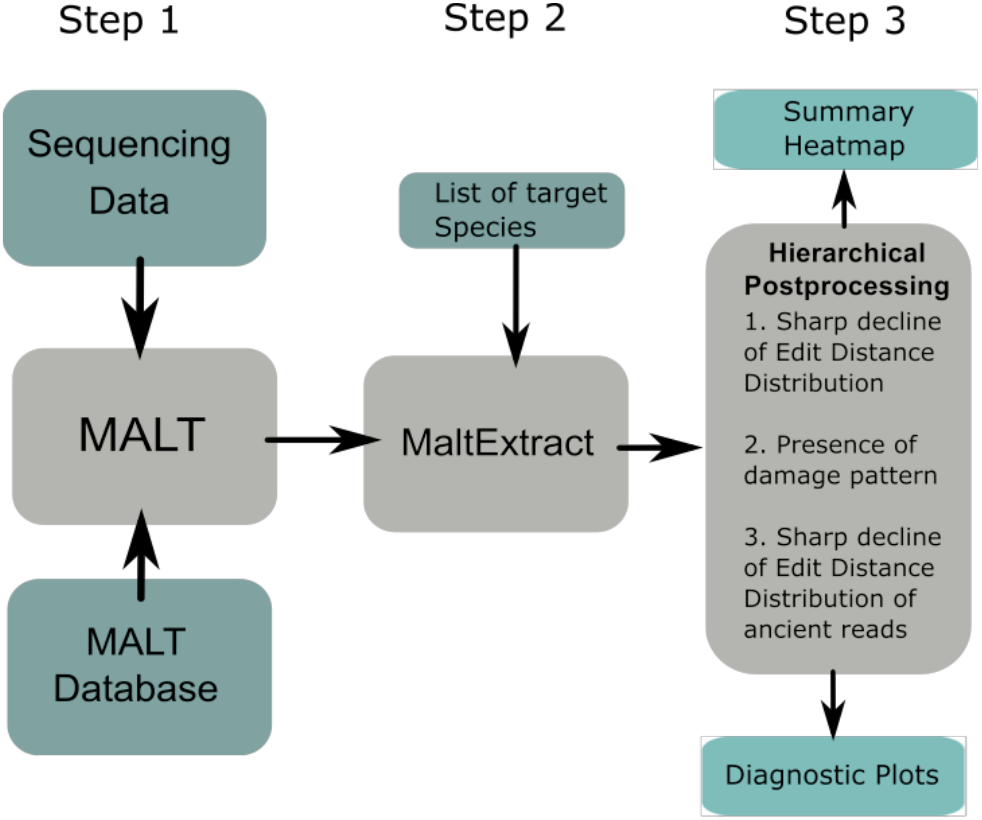
Schematic depiction of HOPS workflow. First, MALT aligns the metagenomic data against its reference database and has an optional mode for processing aDNA reads. MaltExtract then processes the MALT output with various filters and produces various statistics. Finally, post processing procedures provide a comprehensive visualization of the output which can be evaluated to identify putatively positive hits.

#### MALT

MALT (Megan Alignment Tool) (25, 35) is a fast alignment and taxonomic binning tool for metagenomic data that aligns DNA sequencing reads to a user-specified database of reference sequences. Reads are assigned to taxonomic nodes by the naïve Lowest Common Ancestor (LCA) algorithm (36, 37) and are thus assigned to different taxonomic ranks based on their specificity. The default version of MALT is intended for the analysis of metagenomic datasets deriving from modern DNA, and thus it was not designed to accommodate the specific requirements of aDNA analyses. In particular, aDNA damage that manifests as miscoding lesions in sequenced products can lead to an increased number of mismatches, and extensive damage has the potential to prevent alignment or alter taxonomic assignment. Loss of target reads due to DNA damage can hamper species detection as aDNA studies usually begin with shallow sequence data during initial evaluations of sample quality. In addition, archaeological remains often show low DNA yields, and library amplification can result in a high number of PCR duplicates, which can falsely inflate quantitative estimates of taxa.

To account for such shortcomings, we introduce a modified version of MALT that is specifically tailored to the analysis of aDNA data. In this modified version, PCR duplicates are removed by eliminating reads identical to those already aligned. In addition, reads are optionally filtered for a minimum Wootton & Federhen complexity (38) in order to remove low complexity reads. Furthermore, to accommodate aDNA damage during alignment, C>T substitutions are ignored in the first five positions from the 5’-end and G>A substitutions are ignored in first five positions from the 3’-end.

### MaltExtract and post-processing

The core of HOPS is formed by the newly developed MaltExtract module. Without MaltExtract the result files produced by MALT (RMA6 format) can only be evaluated manually with the metagenomic analysis tool MEGAN (39). Such analysis becomes infeasible when working with large data sets, in which each sample must be separately searched for a long list of candidate organisms, a process that is both laborious and prone to user error. MaltExtract provides an automated approach for the assessment of the alignment information stored in RMA files generated by MALT. It automatically retrieves and assesses information on various evaluation criteria for all taxonomic nodes that match a given list of target species.

MaltExtract obtains information on edit distance, read length distribution, coverage distribution and alignment mismatch patterns in order to identify and authenticate the presence of species-specific ancient DNA. Furthermore, MaltExtract allows data filtering for maximum read length, minimum percent identity, minimum complexity, and aDNA damage patterns.

Accuracy in taxonomic read assignment is evaluated in a three-step procedure that includes ancient authentication criteria (Figure 2). The first step evaluates the read assignment to a taxonomic node. Incorrect read assignments can occur when databases are incomplete: many species in a metagenomic sample may have no representative reference genome in the database, and hence become erroneously assigned to the closest genetic match, which could belong to a different species, or even genus. Mapping to an incorrect species generally results in an increased number of mismatches across the read that is evident in the edit distance distribution (Figure 2A). By contrast, if the sequenced reads are assigned to the correct reference species, the edit distance distribution should continuously decline, with most of the reads showing no or only a few mismatches, mostly resulting from aDNA damage or evolutionary divergence of the modern reference from the ancient genome. We summarize the shape of the edit distance distribution by a score we term *negative difference proportion (−Δ%)*, which leverages the difference in sequencing read counts between neighboring mismatch categories (Figure S1). The −Δ% takes values between 0 and 1, where 1 indicates a strictly declining edit distance distribution. While true positives have a −Δ% of 1 when enough endogenous species-specific sequencing reads are present, we use a threshold of −Δ% > 0.9 to account for possible perturbations due to stochasticity in the edit distance distribution when few reads (~10-20) are present. As such, this permits the detection of even low abundant taxa.

**Figure 2.**
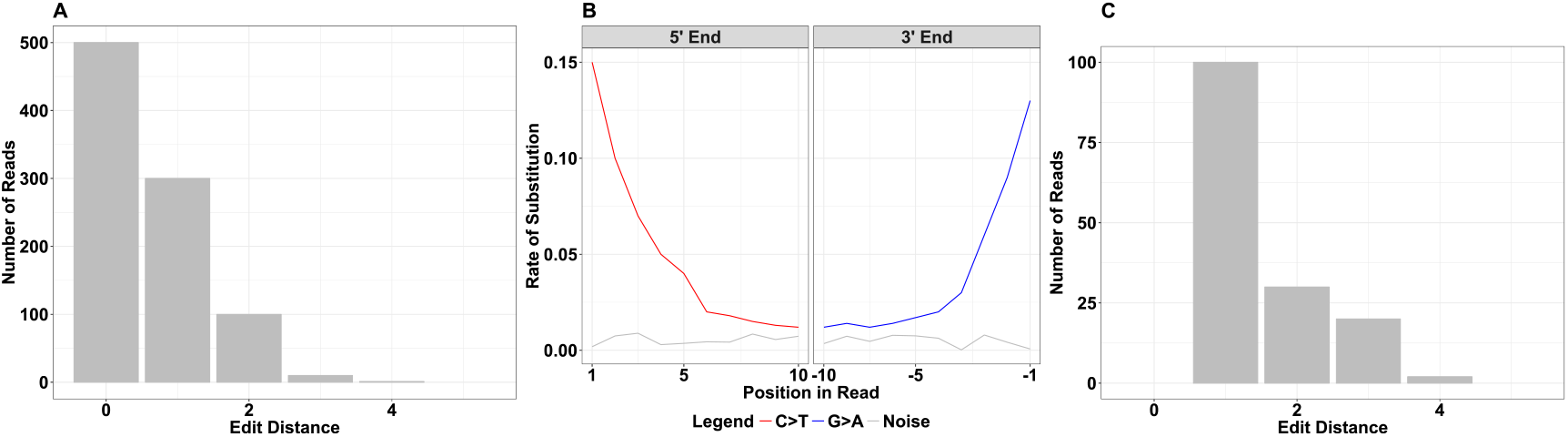
Post-processing steps in HOPS. Three hierarchical post-processing steps are used in HOPS. (A) First, the edit distance distribution is required to show a decline (black lines). (B) Second, the alignments are assessed for C>T and G>A mismatches typical for aDNA; by default, any such damage is considered sufficient. (C) Third, the edit distance distribution of reads showing damage is evaluated.

In a second step, the ancient origin of the DNA is evaluated through analysis of DNA miscoding lesion patterns (Figure 2B). The most prominent modification observed is deamination of cytosine into uracil, which is read as a thymine by the polymerase. This leads to an overrepresentation of C>T substitutions at the 5’ end and correspondingly G>A substitutions at the 3’ end (34, 40). Evaluation of damage patterns is mandatory in any ancient DNA study. MaltExtract reports the rates of substitutions for the leading and trailing 10 positions of the read alignment. The default post-processing settings require only a single miscoding lesion to be present in at least one read to qualify as exhibiting damage. This maximizes sensitivity and allows authentication to function largely independently of read depth.

As a third and final criterion, we evaluate the accuracy of taxonomic assignment for all aligned reads exhibiting aDNA damage. For this we assess again the edit distance distribution using the −Δ% score, but now this is only performed for damaged reads (Figure 2C). In this step, a greater number of assigned reads (>100) is required for reliable edit distance evaluation due to the fact that not all ancient reads are expected to exhibit damage.

The MaltExtract output is saved in a structured output folder with a summary file of the processed input and subfolders for each evaluation criterion. The post-processing tool generates a summary highlighting which of the target species passed one or more evaluation criteria for each sample, as well as detailed diagnostic plots displaying the evaluation criteria for each supported target species (Figure S2).

### Assessment of taxonomic assignment on simulated data

The naïve LCA algorithm (36), which is part of HOPS, assigns reads to different taxonomic levels depending on the specificity of sequence matches. Taxonomic assignment thus depends on the structure of the underlying reference database, and it is critical to understand the expected taxonomic placement of sequenced reads from each microbial pathogen in order to successfully identify them. To analyze the taxonomic placement of a test set of 33 bacterial pathogens and to assess the performance of HOPS we simulated sequencing reads that included artificial DNA damage and spiked them into dentine, dental calculus, bone and soil metagenomic backgrounds (see Table 1).

**Table 1.** Metagenomic backgrounds used for simulated data sets

| ID | Source | Age (Period) | Treatment | Reference |
| --- | --- | --- | --- | --- |
| KT31calc | Calculus | Medieval | No UDG | (50) |
| LP39.10 | Dentine | 2920-2340 BCE | No UDG | (51) |
| MK5.001 | Dentine | 3348-3035 BCE<br>3619-3366 BCE | UDG half | (52) |
| TÖSM_1a | Bone | 6000-5500 BCE | UDG half | (53) |
| Soil | Soil | - | No UDG | (25) |

Applying the HOPS pipeline, we recovered 98% of the simulated reads for 32 of the 33 bacterial taxa of interest (see Figure 3). The one exception was *Mycobacterium avium subspecies paratuberculosis K10* for which 23% of simulated reads were assigned to an incorrect *Mycobacterium avium subspecies paratuberculosis* strain. Our analysis shows that in most cases the taxonomic levels “species” and “complex” (e.g. *Mycobacterium tuberculosis* complex and *Yersinia pseudotuberculosis* complex) correctly accumulate the vast majority of the simulated pathogen reads. However, noteworthy exceptions were *Brucella abortus*, *Brucella melitenis* and *Bordetella pertussis*. Upon further investigation, we found that many species within the genera *Brucella* and *Bordetella* show a high degree of sequence similarity, thus causing the majority of the reads deriving from these pathogens to be assigned at the genus level. By contrast, read assignment was found to be very specific for five taxa (*Treponema denticola* ATCC 35405, *Clostridium tetani* E89, *Clostridium botulinum* E3 str. Alaska E43, *Streptococcus gordonii* str. Challis substr. CH1 and *Clostridium botulinum* BKT015925), resulting in the majority of reads deriving from these taxa to be correctly assigned at the strain level. For *Salmonella enterica* subsp. *enterica* most reads were assigned at the subspecies level. The results of this test provide a guide for the levels of taxonomic identification that should be considered when searching for any of the 33 queried bacterial species in experimental ancient datasets.

**Figure 3.**
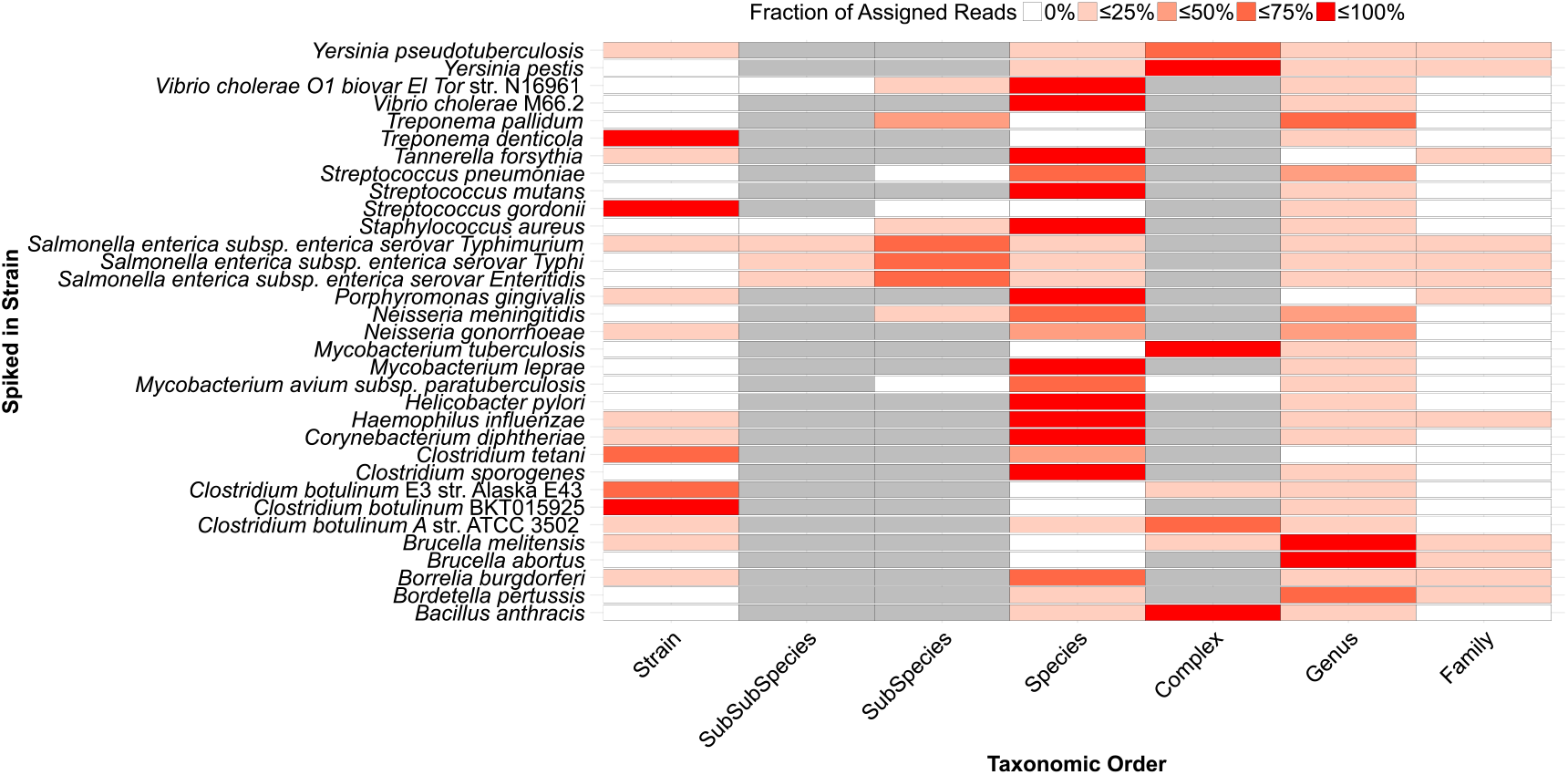
Assignment of simulated reads to taxonomic levels for 33 bacterial pathogens. The fraction of simulated reads (red gradient) per reference (y axis) assigned to a specific node across different levels of the taxonomy (x axis). The levels of taxonomy not defined for a species are shown in grey.

### Optimization of MALT for aDNA

Because MALT was designed for taxonomic binning of modern genetic data, adapting it to be used on aDNA required altering the original MALT implementation to tolerate terminal substitutions consistent with aDNA damage so that they would not interfere with the percent identity filter. To evaluate the efficacy of this modification, we compared the performance of the modified, damage tolerant version of MALT to the default version using simulated *Y. pestis* data with high damage (~40%) and three different percent identity filters: 85%, 95% and 99% (Figure 4).

**Figure 4.**
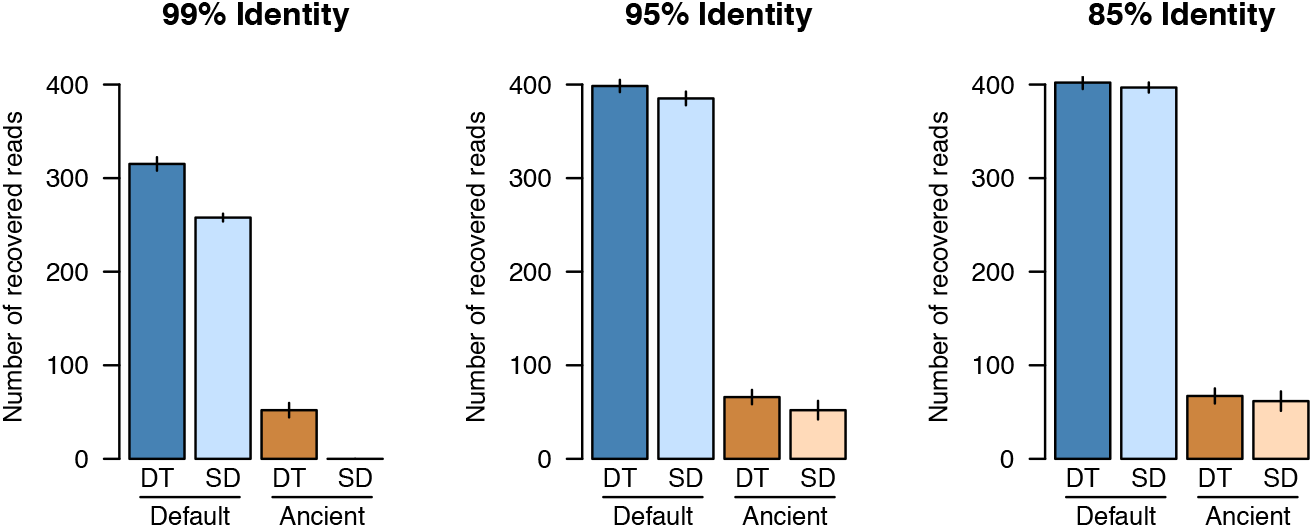
Comparison of the number of successfully recovered *Y. pestis* reads using standard (SD) and damage tolerant (DT) MALT with minimum percent identities of (A) 85%, (B) 95% and (C) 99%. Shown are the recovered reads from the “default” (all reads) and “ancient” (reads with damage) modes in MALT, with the same 500 reads being spiked into the metagenomic backgrounds. Error bars show the standard error of five independent technical replicates for each analysis.

As expected, the greatest difference was observed when applying the stringent 99% identity filter, for which the damage tolerant MALT version recovered ~20% more reads than the standard MALT version. Additionally, only the modified version was able to recover reads with simulated damage under these parameters. At 95% identity, only a small difference could be observed between the two MALT versions, while results were almost identical at an 85% identity level. Taken together, the damage tolerant MALT version provides an advantage when searching for a given pathogen using stringent filtering criteria.

### Performance comparison of HOPS, Kraken and MIDAS on simulated data

We next evaluated the performance of HOPS by comparing it to two commonly used metagenomics profiling tools: MIDAS (30), a marker gene-based taxonomic classifier, and Kraken (31), which performs taxonomic classification based on k-mer matching to a database of complete genomes. The marker gene database of MIDAS lacked representation for *Yersinia pseudotuberculosis, Bordetella pertussis* and *Brucella melitensis*. Therefore, MIDAS could only be evaluated for 30 of the 33 bacterial pathogens in the simulated data sets. For Kraken, we downloaded the bacterial database, which lacked a reference genome to *Clostridium sporogenses*.

HOPS consistently detected all 33 pathogens in all backgrounds and among replicates with as few as 50 reads (see Figure 5A). Kraken failed to identify *Brucella abortus* and *Mycobacterium tuberculosis* in some replicates with only 50 simulated pathogen reads, but otherwise had a sensitivity of 100%; however, it was prone to a high false positive rate (see below). The sensitivity of MIDAS was far lower than for Kraken and HOPS. Even with 5000 simulated pathogen reads for each species, MIDAS detected only 11 of the 30 possible bacterial pathogens. This can be explained by the limited sensitivity of marker gene based approaches, which require relatively high sequencing coverage in order to ensure adequate representation of the marker genes needed for identification. This is further evident as MIDAS’ sensitivity is more heavily influenced by the number of simulated reads than Kraken and HOPS.

**Figure 5.**
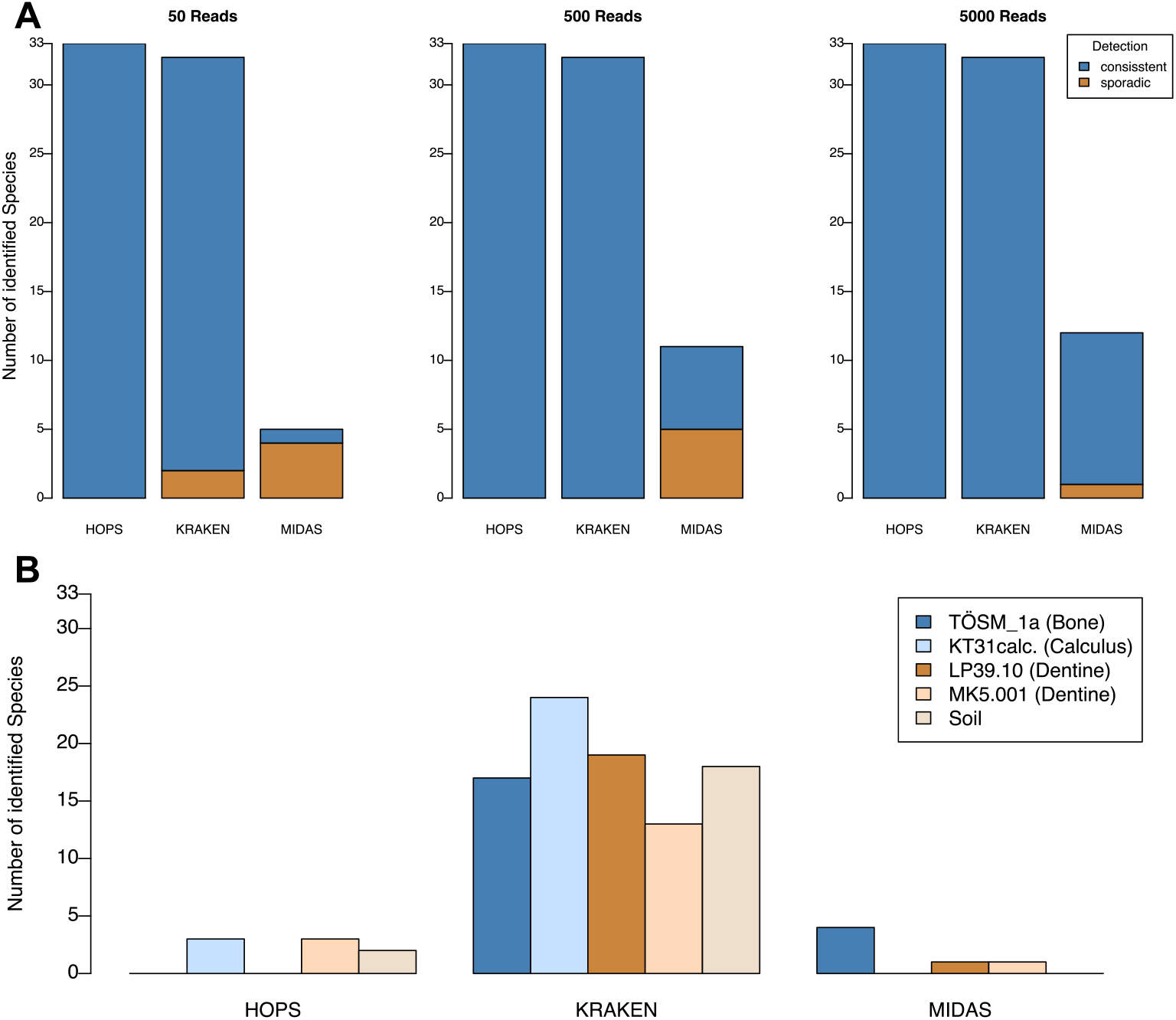
Performance comparison of HOPS, Kraken and MIDAS. (A) HOPS outperforms other tools, successfully and consistently identifying all 33 target bacteria, even when represented by as few as 50 reads. (B) Number of target species identified in the metagenomic background files (negative controls) for HOPS, Kraken and MIDAS.

### Negative controls

To assess false positive assignments, we queried all five metagenomic datasets for detectable signatures of the 33 test bacterial pathogens using HOPS, Kraken and MIDAS in the five metagenomic backgrounds prior to the addition of simulated pathogen reads. Kraken showed the highest susceptibility to false positives (see Figure 5B; Table S1). In this analysis, Kraken detected 24 (73%) pathogens in calculus, 19 (58%) in dentine, 13 (39%) in bone and 18 (55%) in soil. Most problematically, *Mycobacterium tuberculosis* and *Bordetella pertussis* were detected by Kraken in every metagenomic background.

Unexpectedly, MIDAS detected oral streptococci, *Tannerella forsythia, Treponema denticola* and *Porphyromonas gingivalis* in the dentine samples but not in calculus, where they are normally found. Overall, MIDAS produced fewer identifications than Kraken, but such a result is expected given its reliance on marker gene-based detection, which limits identification to only abundant taxa.

HOPS detected four test pathogens in the metagenomic background datasets: *Clostridium tetani* (soil), *Streptococcus mutans* (calculus, dentine), *Treponema denticola* (calculus, dentine), and *Porphyromonas gingivalis* (calculus only). Because *C. tetani* is ubiquitous in soil, and all other detected bacteria are commensals of the human oral cavity, their identification via both MIDAS and HOPS likely reflects true positives. Taken together, HOPS and MIDAS have a lower tendency toward false positive assignments. Kraken’s increased vulnerability for aberrant assignments likely relates to the absence of an alignment step, which is necessary for reliable species evaluation in both modern and ancient contexts.

### Positive Controls

In addition to performing tests using simulated data, we also tested HOPS, Kraken and MIDAS on 25 ancient metagenomic datasets known to be positive for bacterial pathogens (Table 2). They consisted of both shotgun and capture data and they varied in sequencing depth in accordance with experimental conditions and method of data generation.

**Table 2.** Metagenomic samples used as positive controls

| ID | Reconstructed Bacteria | Sequencing reads | Data type | Reference |
| --- | --- | --- | --- | --- |
| 10C | *Salmonella enterica* | 1,017,400 | Shotgun | (25) |
| 35C | *Salmonella enterica* | 986,908 | Shotgun | (25) |
| RK1001.C0101 | *Yersinia pestis* | 7,023,370 | Shotgun | (17) |
| GEN_72 | *Yersinia pestis* | 7,663,408 | Shotgun | (17) |
| 549_O | *Yersinia pestis* | 1,520,471 | Shotgun | (16) |
| JK3031UDG | *Yersinia pestis* | 4,059,016 | Shotgun (UDG) | (16) |
| JK2370UDG | *Yersinia pestis* | 52,858,027 | Shotgun (UDG) | (16) |
| RT6 | *Yersinia pestis* | 6,706,316 | Shotgun (UDG) | (18) |
| 1343UnTal85 | *Yersinia pestis* | 3,462,216 | Shotgun | (17) |
| 6Post | *Yersinia pestis* | 2,546,695 | Shotgun | (17) |
| KunilaII | *Yersinia pestis* | 1,007,417 | Shotgun | (17) |
| RISE00 | *Yersinia pestis* | 6,000,000 | Shotgun | (13) |
| RISE139 | *Yersinia pestis* | 6,000,000 | Shotgun | (13) |
| RISE386 | *Yersinia pestis* | 6,000,000 | Shotgun | (13) |
| RISE397 | *Yersinia pestis* | 6,000,000 | Shotgun | (13) |
| RISE505 | *Yersinia pestis* | 6,000,000 | Shotgun | (13) |
| RISE509 | *Yersinia pestis* | 6,000,000 | Shotgun | (13) |
| RISE511 | *Yersinia pestis* | 6,000,000 | Shotgun | (13) |
| 54 | *Mycobacterium tuberculosis* | 70,897 | Shotgun | (21) |
| 58 | *Mycobacterium tuberculosis* | 114,555 | Shotgun | (21) |
| 64 | *Mycobacterium tuberculosis* | 160,310 | Shotgun | (21) |
| 54 | *Mycobacterium tuberculosis* | 5,000,000 | Capture (UDG) | (21) |
| 58 | *Mycobacterium tuberculosis* | 5,000,000 | Capture (UDG) | (21) |
| 64 | *Mycobacterium tuberculosis* | 5,000,000 | Capture (UDG) | (21) |
| P1P2 | *Helicobacter pylori* | 5,000,000 | Capture (UDG) | (27) |

HOPS and Kraken share 100% sensitivity for the detection of target bacterial pathogens in every sample. By contrast, MIDAS only detected the correct bacterial pathogen in 22 out of 25 samples. Again, MIDAS sensitivity was likely reduced due to the marker gene-based approach. These results highlight the advantage of whole-genome based approaches like MALT and Kraken that take advantage of every sequenced read.

### Runtimes

To calculate the runtime for each program, we used simulated metagenomic files that each contained five million sequencing reads (see Methods). For each file, HOPS required an average of 3307±820 seconds for the MALT step, 16±1 seconds for the MaltExtract step and 1±0 seconds for post processing, for a total of approximately 55 minutes of analysis time per file. Kraken took on average 72±16 seconds to run *Kraken_alignment* and 22±3 for *Kraken_translate*, a total of 1.5 minutes, and the MIDAS pipeline processed each file in an average of 4±1 seconds. HOPS by far required the highest runtimes of the three tools, but most of this time was required for sequence alignment, a step that, although time consuming, increases detection sensitivity, reduces false positives, and enables the authentication of aDNA reads.

## Discussion

The field of archaeogenetics faces several challenges, such as the low amount of endogenous target DNA, the highly degraded nature of the DNA, and an unknown and diverse metagenomic background signal that accumulates during decomposition and centuries spent in a depositional environment. This makes reliable identification and authentication of genuine ancient DNA challenging, particularly when targeting bacterial DNA that is usually only present in small amounts. Furthermore, many bacterial pathogens have close relatives in soil, which necessitates meticulous care when making pathogen identifications.

HOPS provides an automated pipeline for high-throughput ancient bacterial species detection and authentication from metagenomic sequencing data. We compare HOPS to Kraken and MIDAS, two widely used methods for estimating both the presence and abundance of bacterial taxa in metagenomic data. These tools, however, have limited application to the specific challenges of aDNA in terms of degradation and chemical modifications manifest as miscoding lesions. Our analyses highlight the need for a pathogen identification pipeline that accommodates qualities of aDNA data and includes an essential and robust authentication for all ancient read assignments. HOPS provides a fast, reliable, and user-friendly solution to these established limitations.

HOPS was tested on simulated ancient pathogen DNA reads, and it successfully detected all targeted species spiked into metagenomic backgrounds with as few as 50 pathogen reads, representing less than 0.00001 % of the total dataset. In this context, our modified version of MALT, which tolerates mismatches resulting from DNA degradation, prevents a decrease in sensitivity even in cases of heavily damaged aDNA. We demonstrate that the marker gene-based metagenomic profiling tool MIDAS had a much lower sensitivity for pathogen detection compared to HOPS, especially for low coverage data, which is typical of ancient DNA screening datasets. Although the sensitivity of Kraken was similar to HOPS, and while Kraken’s k-mer matching is considerably faster than the precise alignments used in HOPS, Kraken is incapable of validating species assignment and aDNA authenticity, and thus has a lower specificity. This is most clearly demonstrated by our analysis of a metagenomic soil sample in which Kraken detected numerous false positives, including *Mycobacterium tuberculosis* and *Bordetella pertussis* (whooping cough). This is likely due to many soil dwelling bacteria that harbor genetic similarities to these pathogens, such as diverse mycobacerial species and *B. petrii*, a close relative to *B. pertussis* that is a common constituent of environmental datasets. These effects are further compounded by the fact that many environmental microbes have not been genomically characterised and are not part of any reference database, which only increases the potential of false assignments to well-sequenced pathogens. The alignment-based validation procedure implemented in HOPS minimises such false positive assignments, and thus offers greater accuracy in pathogen identification during screening when environmental backgrounds comprise the dominant molecular signal.

A previously published pipeline for the assessment of metagenomic data in archaeogenetics is metaBIT (41). It implements a variety of methods for the detailed assessment of metagenomic composition, which also includes validation of aDNA damage patterns. As metaBIT is based on MetaPhlAn (42), which employs a marker gene based approach in the initial detection step similar to MIDAS, pathogens in low abundance could be missed in its initial steps when applied to shallow sequencing data. An integrated approach combining HOPS and metaBIT might be a promising future strategy for a detailed characterization of microbiomes while at the same time proving a high level of sensitivity for the detection of pathogens. In particular, the analysis of ancient samples that preserve their original microbiome signature, such as dental calculus (43) or coprolites (44) would benefit from a combined application of both methodologies, by using metaBIT to assess the microbial make up and using HOPS for more in depth species authentication.

For all taxonomic classifiers, correct assignment of metagenomic reads is strongly dependent on the quality of the underlying reference sequences. Currently we use a curated database for MALT that contains completed reference sequences and assemblies for bacteria from RefSeq (December 2016). Database sizes are constantly increasing, but much of this growth derives from the addition of redundant sequence data from model organisms, which also creates biases. In this context, methodologies such as SPARSE (45) aim to mitigate against database redundancy by hierarchically structuring reference sequences, which could be employed to further improve HOPS’ specificity and runtime.

In addition, analysis of our simulated dataset allowed for insights into the taxonomic structure of each of the bacterial pathogens in our target list. It became apparent that for some targets the taxonomic species level is not sufficient for identification. This applies to historically important pathogens such as *Y. pestis* or *M. tuberculosis*. Here, evaluation of a higher taxonomic level such as complex is more reliable, while in the case of *Salmonella typhi* (typhoid fever) a lower level (subspecies) is favorable. Therefore, our simulations provide a valuable resource for the optimization of pathogen screening approaches in general.

Here, HOPS was evaluated for its success in screening for bacterial pathogens. Because the reference database is user defined and can amended to include, for example, the NCBI full nucleotide collection (46) or hand-curated sets of reference genomes, tremendous flexibility exists in molecular detection, which could extend to viruses, fungi, and eukaryotes.

## Conclusions

We present a fast, reliable, and user-friendly computational pathogen screening pipeline for ancient DNA that has the flexibility of handling large datasets. HOPS successfully identifies both simulated and actual ancient pathogen DNA within complex metagenomic datasets, exhibiting a higher sensitivity than MIDAS and with fewer false positives than Kraken. HOPS provides a high level of automatization that allows for the screening of thousands of datasets with very little hands-on time, and it offers detailed visualizations and statistics at each evaluation step, enabling a high level of quality control and analytical transparency. HOPS is a powerful tool for high-throughput pathogen screening in large-scale archaeogenetic studies, producing reliable and reproducible results even from remains with exceptionally low levels of pathogen DNA. Such qualities make HOPS a valuable tool for pathogen detection in the rapidly growing field of archaogenetics.

## Methods

### Implementation of MaltExtract

MaltExtract is implemented in Java. It integrates parts of MEGAN’s (39) source code for accessing the RMA file structure and functions from *forester* (https://github.com/cmzmasek/forester) for traversing the taxonomic tree.

### Simulating data to analyse read assignment using the MALT LCA algorithm

Depending on the database structure and sequence similarity between reference sequences, the naïve LCA (36) algorithm will assign reads to different taxonomic units. To inquire how reads are assigned to the taxonomic tree for 33 bacterial pathogens (Table S2), we simulated ancient pathogen DNA reads using gargammel (47) and spiked them into five ancient metagenomic background datasets obtained from bone, dentine, dental calculus and soil (Table 1). The simulated reads carry a unique identifier in their header in order to differentiate them from metagenomic background sequences, which exhibit either full damage patterns or attenuated damage patterns following UDG-half treatment (48). To simulate aDNA damage in the pathogen sequences, we applied damage profiles obtained from previously published ancient *Yersinia pestis* genomes with (13) and without UDG-half (18) treatment. Simulated reads were processed with EAGER (49) and spiked into the metagenomic backgrounds in different amounts (50, 500 or 5000 reads). For each metagenomic background, a typical screening sequencing depth of five million reads were used.

### Evaluation of the damage tolerant version of MALT

To preserve damage patterns when mapping reads with MALT, we modified the source code and compared the performance of the modified and default versions. We therefore created with gargammel (47) test samples that show twice the amount of damage (~40%) usually found in ancient samples (13). Here, we compare both MALT versions for the bacterial pathogen *Yersinia pestis* (CO92 reference). Both versions of MALT were tested with 85%, 95% and 99% minimum percent identity filtering, to investigate the effects of percent identity filtering on the read alignment of aDNA reads.

### Comparison of HOPS to Kraken and MIDAS

HOPS was compared to two metagenomic taxonomic classification tools: Kraken (31) and MIDAS (30). We only executed the first step of MIDAS that matches reads to the marker gene database to determine species abundance. This step was executed on 32 cores with default parameters. The first step is sufficient, as any species undetected in this step would not be detected in the remaining ones. Kraken was set to use 32 cores to align the sample data against its reference database with the preload parameter to load the entire database into memory before starting k-mer alignment. In a second step kraken-translate was executed to transform taxonomy ids into proper species names. For Kraken and MIDAS, we judged a pathogen as correctly identified if at least one read matches to a strain of the correct species to account for the differences in the database contents, methodologies and output formats.

### Databases

In our study, HOPS uses a database containing all complete prokaryotic reference genomes obtained from NCBI (December 1st 2016) with entries containing ‘multi’ and ‘uncultured’ removed (13 entries). In total, 6,249 reference genomes are included in the database. For Kraken we downloaded the bacterial database with Kraken’s kraken-build script (June 1 2017). The Kraken database contains no strain references for *Clostridium sporogenses*. Otherwise it contains at least one reference for all of the simulated bacterial pathogens (Table S2). For MIDAS we used the default reference database (May 24 2016), which contained no representation of *Yersinia pseudotuberculosis, Bordetella pertussis* and *Brucella melitensis*.

### Positive controls

We compare the sensitivity and specificity of HOPS, MIDAS and Kraken using 25 metagenomic datasets previously shown to be positive for one of four microbial pathogens: *Yersinia pestis, Mycobacterium tuberculosis, Salmonella enterica* and *Heliobacter pylori* (Table 2). These positive control samples represent real metagenomic data and therefore contain an unknown number of modern species in addition to the actual recovered bacterial pathogen. Read counts across all samples ranged from 70,897 to 7,000,000 reads. While most datasets were generated by shotgun library screening, four datasets were enriched for pathogen DNA prior to sequencing using DNA capture methods. For all captured datasets and a subset of shotgun datasets, DNA was treated with UDG prior to library construction to remove DNA damage. Both types of datasets were included to evaluate the performance of HOPS on samples with different levels of DNA damage and pathogen abundance.

### Runtimes

To calculate the runtimes for HOPS, Kraken and MIDAS, we used a subset of the simulated files. The subset consisted of all metagenomic background datasets spiked with 5000 reads without technical replicates resulting in a total of 330 metagenomic files. A total of 64 cores and 700 GB of RAM were allocated to each program.

## Declarations

### Availability of data and material

The complete source code of HOPS is available from GitHub (https://github.com/rhuebler/HOPS).

### Authors’ contributions

RH, FMK and AH conceived the study. RH, FMK, CW, KIB, JK and AH designed experiments. RH and FMK implemented software. RH, FMK and AH performed analyses. RH, FMK and AH wrote the manuscript with contributions from all coauthors.

## Acknowledgements

We thank the Department of Archaeogenetics of the Max Planck Institute for the Science of Human History and Julian Susat for beta testing and helpful discussions.

## Supplementary Information

**Figure S1.**
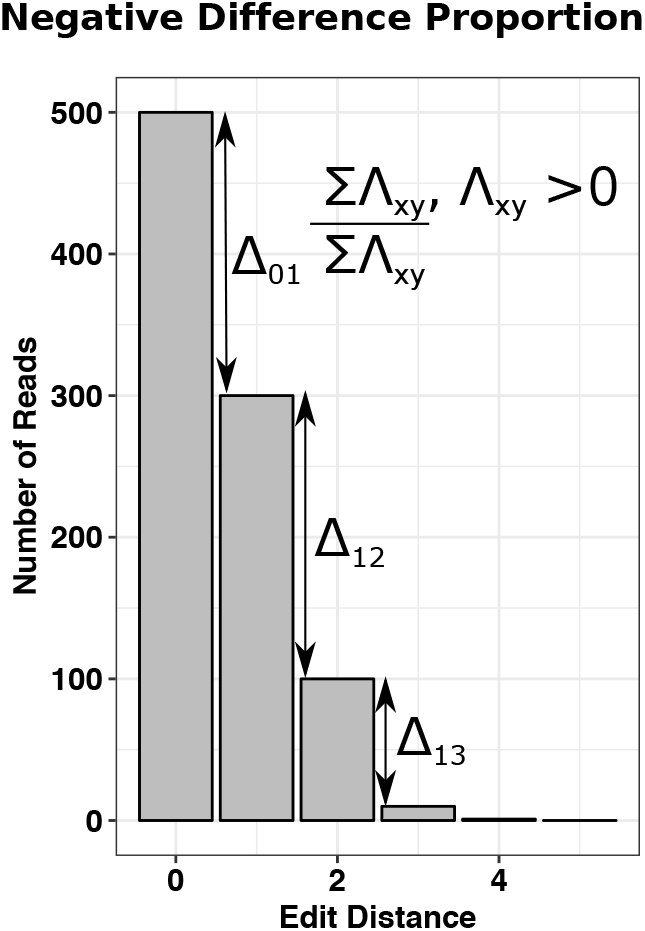
The first and third steps in the HOPS postprocessing protocol require a decline in the Edit Distance distribution.

**Figure S2.**
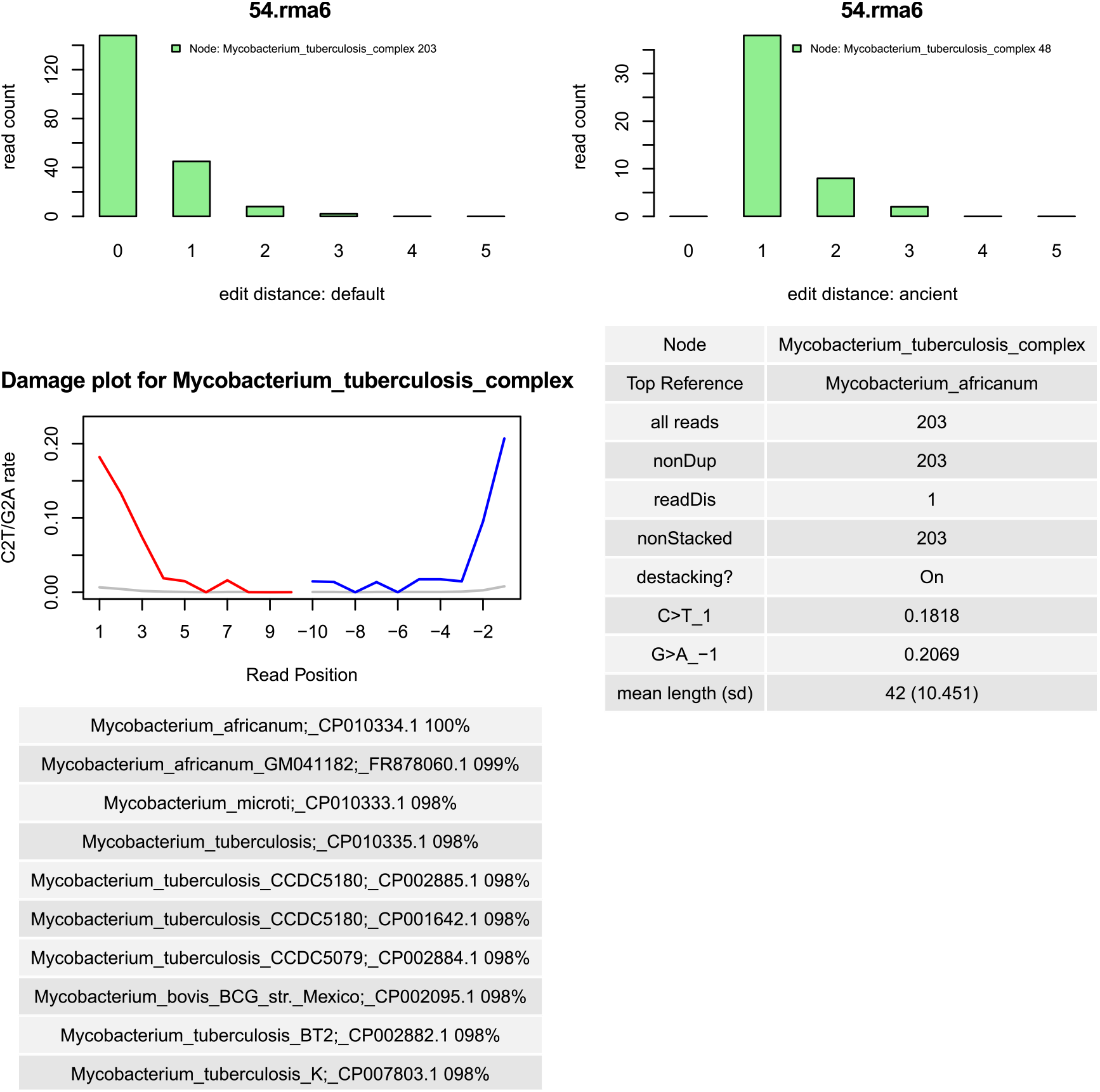
HOPS summary output for a tuberculosis positive sample. Upper left: Edit distance distribution for all reads assigned to M. tuberculosis. Upper right: Edit distance distribution for assigned reads that show a possible DNA damage signal. Middle left: DNA damage plot for assigned reads. Lower left: Top ten references with percentage of aligned reads. Middle right: Summary statistics for assigned reads.

**Table S1.** Results for negative controls. For HOPS the step in the post processing that was reached for each species is indicated (0: No detection; 1: detected with good edit distance distribution; 2: additionally indication for damage; 3: additionally good edit distance distribution for damaged reads). For Kraken the number of k-mers assigned to the species and for MIDAS the number of assigned reads is listed.

| Species | bone_TOSM1a | calculus_KT31calc | dentine_MK5.001 | dentine_LP39.10 | soil | Software |
| --- | --- | --- | --- | --- | --- | --- |
| Bacillus anthracis | 0 | 0 | 0 | 0 | 0 | HOPS |
| Bordetella pertussis | 0 | 0 | 0 | 0 | 0 | HOPS |
| Borrelia | 0 | 0 | 0 | 0 | 0 | HOPS |
| burgdorferi B31 |  |  |  |  |  |  |
| Brucella abortus | 0 | 0 | 0 | 0 | 0 | HOPS |
| Brucella melitensis | 0 | 0 | 0 | 0 | 0 | HOPS |
| Clostridium botulinum | 0 | 0 | 0 | 0 | 1 | HOPS |
| Clostridium sporogenes | 0 | 0 | 0 | 0 | 0 | HOPS |
| Clostridium tetani | 0 | 0 | 0 | 0 | 1 | HOPS |
| Corynebacterium diphtheriae | 0 | 0 | 0 | 0 | 0 | HOPS |
| Haemophilus influenzae | 0 | 0 | 0 | 0 | 0 | HOPS |
| Helicobacter pylori | 0 | 0 | 0 | 0 | 0 | HOPS |
| Mycobacterium avium subsp. paratuberculosis | 0 | 0 | 0 | 0 | 0 | HOPS |
| Mycobacterium leprae | 0 | 0 | 0 | 0 | 0 | HOPS |
| Mycobacterium tuberculosis | 0 | 0 | 0 | 0 | 0 | HOPS |
| Neisseria gonorrhoeae | 0 | 0 | 0 | 0 | 0 | HOPS |
| Neisseria meningitidis | 0 | 0 | 0 | 0 | 0 | HOPS |
| Porphyromonas gingivalis | 0 | 3 | 0 | 0 | 0 | HOPS |
| Salmonella enterica subsp. enterica | 0 | 0 | 0 | 0 | 0 | HOPS |
| Staphylococcus aureus subsp. aureus | 0 | 0 | 0 | 0 | 0 | HOPS |
| Streptococcus gordonii str. | 0 | 0 | 0 | 1 | 0 | HOPS |
| Streptococcus mutans | 0 | 3 | 0 | 2 | 0 | HOPS |
| Streptococcus pneumoniae | 0 | 2 | 0 | 0 | 0 | HOPS |
| Tannerella forsythia | 0 | 0 | 0 | 0 | 0 | HOPS |
| Treponema denticola | 0 | 0 | 0 | 1 | 0 | HOPS |
| Treponema pallidum subsp. pallidum | 0 | 0 | 0 | 0 | 0 | HOPS |
| Vibrio cholerae | 0 | 0 | 0 | 0 | 0 | HOPS |
| Yersinia pestis | 0 | 0 | 0 | 0 | 0 | HOPS |
| Yersinia pseudotuberculosis | 0 | 0 | 0 | 0 | 0 | HOPS |
|  | bone_TOSM1a | calculus_KT31calc | dentine_MK5.001 | dentine_LP39.10 | soil | Kraken |
| Bacillus anthracis | 0 | 5 | 1 | 0 | 0 | Kraken |
| Bordetella pertussis | 1 | 12 | 20 | 4 | 10 | Kraken |
| Borrelia burgdorferi B31 | 0 | 0 | 0 | 0 | 0 | Kraken |
| Brucella abortus | 0 | 0 | 0 | 0 | 1 | Kraken |
| Brucella melitensis | 0 | 4 | 0 | 0 | 1 | Kraken |
| Clostridium botulinum | 40 | 182 | 0 | 21 | 40 | Kraken |
| Clostridium sporogenes | 0 | 0 | 0 | 0 | 0 | Kraken |
| Clostridium tetani | 13 | 50 | 1 | 2 | 240 | Kraken |
| Corynebacterium diphtheriae | 4 | 79 | 9 | 13 | 11 | Kraken |
| Haemophilus influenzae | 2 | 1617 | 0 | 2 | 3 | Kraken |
| Helicobacter pylori | 8 | 12 | 0 | 0 | 4 | Kraken |
| Mycobacterium avium subsp. paratuberculosis | 13 | 8 | 39 | 6 | 26 | Kraken |
| Mycobacterium leprae | 16 | 19 | 22 | 13 | 21 | Kraken |
| Mycobacterium tuberculosis | 53 | 46 | 208 | 55 | 117 | Kraken |
| Neisseria gonorrhoeae | 4 | 2407 | 2 | 3 | 8 | Kraken |
| Neisseria meningitidis | 4 | 4448 | 0 | 9 | 12 | Kraken |
| Porphyromonas gingivalis | 2 | 13925 | 0 | 6 | 0 | Kraken |
| Salmonella enterica subsp. enterica | 8 | 54 | 16 | 24 | 16 | Kraken |
| Staphylococcus aureus subsp. aureus | 1 | 19 | 2 | 4 | 4 | Kraken |
| Streptococcus gordonii | 0 | 71751 | 0 | 15 | 0 | Kraken |
| Streptococcus mutans | 0 | 401 | 0 | 17 | 0 | Kraken |
| Streptococcus pneumoniae | 0 | 3224 | 0 | 0 | 0 | Kraken |
| Tannerella forsythia | 8 | 31294 | 14 | 18 | 14 | Kraken |
| Treponema denticola | 0 | 25264 | 0 | 4 | 0 | Kraken |
| Treponema pallidum subsp. pallidum | 0 | 0 | 0 | 0 | 0 | Kraken |
| Vibrio cholerae | 6 | 60 | 1 | 6 | 5 | Kraken |
| Yersinia pestis | 0 | 4 | 0 | 0 | 0 | Kraken |
| Yersinia pseudotuberculosis | 15 | 21 | 7 | 39 | 7 | Kraken |
|  | bone_TOSM1a | calculus_KT31calc | dentine_MK5.001 | dentine_LP39.10 | soil |  |
| Bacillus anthracis | 0 | 0 | 0 | 0 | 0 | Midas |
| Bordetella pertussis | 0 | 0 | 0 | 0 | 0 | Midas |
| Borrelia burgdorferi B31 | 0 | 0 | 0 | 0 | 0 | Midas |
| Brucella abortus | 0 | 0 | 0 | 0 | 0 | Midas |
| Brucella melitensis | 0 | 0 | 0 | 0 | 0 | Midas |
| Clostridium botulinum | 0 | 0 | 0 | 0 | 0 | Midas |
| Clostridium sporogenes | 0 | 0 | 0 | 0 | 0 | Midas |
| Clostridium tetani | 0 | 0 | 0 | 0 | 0 | Midas |
| Corynebacterium diphtheriae | 0 | 0 | 0 | 0 | 0 | Midas |
| Haemophilus influenzae | 0 | 0 | 0 | 0 | 0 | Midas |
| Helicobacter pylori | 0 | 0 | 0 | 0 | 0 | Midas |
| Mycobacterium avium subsp. paratuberculosis | 0 | 0 | 0 | 0 | 0 | Midas |
| Mycobacterium leprae | 0 | 0 | 0 | 0 | 0 | Midas |
| Mycobacterium tuberculosis | 0 | 0 | 0 | 0 | 0 | Midas |
| Neisseria gonorrhoeae | 0 | 0 | 0 | 0 | 0 | Midas |
| Neisseria meningitidis | 0 | 0 | 0 | 0 | 0 | Midas |
| Porphyromonas gingivalis | 0 | 0 | 0 | 29 | 0 | Midas |
| Salmonella enterica subsp. enterica | 0 | 0 | 0 | 0 | 0 | Midas |
| Staphylococcus aureus subsp. aureus | 0 | 0 | 0 | 0 | 0 | Midas |
| Streptococcus gordonii str. | 0 | 0 | 0 | 0 | 0 | Midas |
| Streptococcus mutans | 0 | 0 | 0 | 0 | 0 | Midas |
| Streptococcus pneumoniae | 0 | 0 | 0 | 1 | 0 | Midas |
| Tannerella forsythia | 1 | 0 | 1 | 26 | 0 | Midas |
| Treponema denticola | 0 | 0 | 0 | 42 | 0 | Midas |
| Treponema pallidum subsp. pallidum | 0 | 0 | 0 | 0 | 0 | Midas |
| Vibrio cholerae | 0 | 0 | 0 | 0 | 0 | Midas |
| Yersinia pestis | 0 | 0 | 0 | 0 | 0 | Midas |
| Yersinia pseudotuberculosis | 0 | 0 | 0 | 0 | 0 | Midas |

**Table S2.**
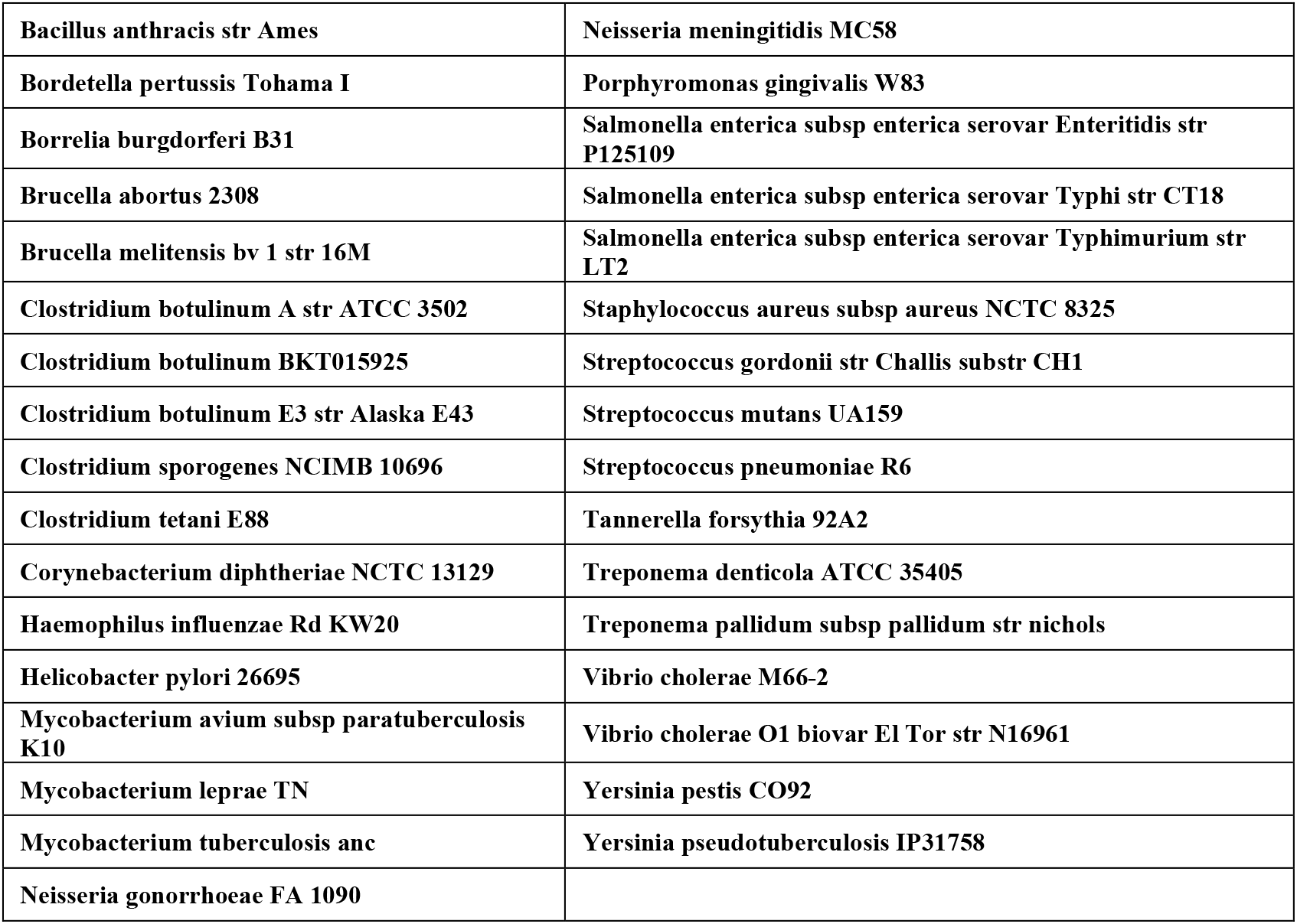
Genomes used to generate simulated ancient pathogen DNA data sets

